# Two-photon voltage imaging of spontaneous activity from multiple neurons reveals network activity in brain tissue

**DOI:** 10.1101/2020.01.29.926014

**Authors:** Binglun Li, Mariya Chavarha, Yuho Kobayashi, Satoshi Yoshinaga, Kazunori Nakajima, Michael Z. Lin, Takafumi Inoue

## Abstract

Recording the electrical activity of multiple neurons simultaneously would greatly facilitate studies on the structure and function of neuronal circuits. Using fluorescent genetically encoded voltage indicators (GEVI) would be especially desirable, as it would allow cell type-selectivity, longitudinal recordings, and further optical manipulations. By expressing the GEVI ASAP3 via *in utero* electroporation and rapidly imaging neurons in densely labelled tissues via random-access multi-photon microscopy, we achieve voltage recording of multiple neurons in brain slice with single-trial single-voxel resolution. This approach enables monitoring of subthreshold membrane potential changes and action potentials from multiple locations in soma and dendrites for tens of minutes. By optically recording spontaneous electrical activities in somatosensory cortex neurons, we provide evidence for the development of intralaminar horizontal connections in layer 2/3 with greater sensitivity than calcium imaging. Single-trial optical voltage recordings using ASAP3 thus enables the investigation of network connectivity at cellular resolution.

## Introduction

The precise recording of membrane voltage in multiple neurons is critical for understanding how the activity of neuronal circuits represent and process information, and thereby underlie brain function. Traditionally measurements of neuronal activity have been performed using electrophysiology, which allows direct and accurate readout of voltage dynamics. Subsequently developed optical methods using fluorescent calcium indicators have enabled easier and less invasive recordings with better spatial resolution and larger target area^1, 2^. Over the last decade, optical imaging with genetically encoded calcium indicators (GECis) such as GCaMPs has become the most prominent approach, enabling studies in sparsely labelled populations or in specific cell types^3–5^.

While imaging calcium demonstrates the advantages of optical imaging of neuronal activity, calcium is only a proxy of electrical activity and its dynamics are significantly slow. In contrast to GECis, genetically encoded voltage indicators (GEVIs) can directly report neuronal electrical activity, but voltage imaging is extremely challenging compared with calcium imaging due to several intrinsic constraints. First, as voltage indicators must reside within the plasma membrane to sense changes in transmembrane voltage^6^, the number of responding indicator molecules, and thereby changes in photon flux per voltage transient are significantly limited ^7^. In addition to the unfavorable spatial constraint on voltage indicators, the transient nature of voltage also poses a problem. While GECis can be sampled at slow frame rates of ~30 Hz to detect the long-lasting calcium fluctuations that follow action potentials, sub-millisecond sampling rates are required to detect the transient optical spikes of fast GEVIs in response to action potentials^7^. The higher sampling rate resulting fewer photons collected per frame leads to lower signal-to-noise ratios (SNRs). Excitation power can be increased to boost SNR, but at the price of greater photobleaching and phototoxicity^8^. Given these constraints, to detect fast voltage transients by voltage imaging, GEVIs must have high response amplitudes and fast response kinetics, and be bright and photostable. At the same time, methods to measure GEVI signals must maximize photon flux from the limited area of plasma membrane over the short durations of voltage transients, i.e. be both sensitive and fast.

After years of intense engineering efforts, the latest generation of GEVIs now have sufficiently improved performance to be deployed in functional *in vivo* studies^7, 9, 10^. For this study we selected a recently reported ASAP3 for its fast kinetics (fast time constant of 0.94 ± 0.06 ms at physiological temperatures) and high responsivity (17.0 ± 0.8% per action potential (AP) in cultured neurons). Most importantly, ASAP3 maintains its responsivity to voltage changes under two-photon excitation, which enabled discrimination of individual spikes in fast-firing neurons in tissue ^7^.

Although GEVIs have been used to monitor electrical activity in neuronal tissue using widefield single-photon microscopy^5, 9, 11, 12^, two-photon microscopy is a better choice in scattering tissues as it permits deep tissue penetration with low background fluorescence and low phototoxicity^13^. With high labeling densities, two-photon excitation is especially useful as individual neurons can be excited without exciting all other neurons along the light path, whereas these other neurons would contribute to background in 1-photon imaging. However, traditional two-photon imaging involves laser scanning at low sampling rates and miniscule dwell times (e.g. 30-Hz sampling of a 512x 512 frame translates to visiting a given location for only 0.1 µs per 33 ms interval). While sufficient for imaging GECI responses, which last for > 100 ms, these speeds are inadequate for detecting GEVI spikes, which last only several milliseconds for the most responsive indicators.^7, 14^ Several strategies have been developed to achieve high sampling rates required for voltage imaging. Measurements performed from a single spot or a line scan across cell membrane have been used to monitor neuronal activity on millisecond-timescales, but recordings are limited to a single or several cells that can be transected by a line^2^. An alternative strategy, random-access multi-photon (RAMP) microscopy ^15, 16^ relies on acousto-optic deflector (AOD) to instantaneously redirect the laser beam among preselected voxels of any location. The high sampling rates achievable with RAMP enabled monitoring of fast voltage signals at sub-millisecond temporal resolution with chemical voltage indicators in cultured neurons^17^, in acute slice^18^, and *in vivo*^7^. In this study, we took advantage of the high sampling rate and random-access scanning to record membrane potential from multiple neurons in brain slices to investigate neuronal circuits in acute brain slices of murine cerebral cortex.

Recording voltage signals from multiple neurons requires a labeling strategy to achieve strong but sparse expression of a GEVI in neurons. A high density of voltage indicator molecules in plasma membrane is crucial for achieving high SNR, while sparse labeling is useful to avoid contact between the plasma membranes of GEVI-expressing cells^8^. In utero electroporation (IUE) allows transfection in specific cell populations depending on the timing and targeted region of electroporation^19^ Conveniently, IUE usually results in a relatively sparser labeling than viral delivery^1^ Also, as GEVI expression begins at prenatal stages, IUE allows monitoring of neuronal activity during development. Thus, we chose IUE to express ASAP3 in specific layers of murine cerebral cortex.

As “cells that fire together wire together”, synchronous firing is the most basic temporal relationship in neural circuits ^20^. Precise spike synchrony is commonly observed in many brain areas^21, 22^. It plays a crucial role in encoding and decoding of neuronal language ^23^, and could reveal the connectivity of the underlying network^24^. Multi-electrode array^25^, multi-cell patch clamping^26^, and calcium imaging.^4, 27^ have been used to detect spike synchrony in local neuron connections. In this study, we investigated the neuron connection with optical voltage recording by combining ASAP3, RAMP and IUE. It is a more efficient and easier approach than electrical measurements, providing better time resolution than calcium imaging.

As a demonstration, we used this approach to study neural pairwise synchronization in mouse cerebral cortex. In mammalian cortex, neurons in layer (L) 2/3 have functional intralaminar horizontal connection and also connect with L4 neurons^28^ but development of these neuronal connections has not been fully understood at cellular resolution. Previous studies using electrophysiology ^29^, photostimulation ^30^ and calcium imaging ^31^ demonstrated a negative correlation between horizontal distance of neuron pair and their correlation strength in L2/3 in adult mice, but a different study reported that network activity in L2/3 was highly synchronous during the first postnatal week and unsynchronized after the second postnatal week^27^. Thus, the developmental changes in horizontal correlation in L2/3 of somatosensory cortex remain to be clarified. In this study, we monitored the spontaneous activity of L2/3 neurons in somatosensory cortex and investigated the spatial/functional relation between neurons during postnatal development, taking advantages of the high brightness and high sensitivity of ASAP3, relatively sparse but strong expression with IUE, and the high speed of RAMP.

## Results

Following ASAP3 gene delivery to murine cerebral cortex via IUE (Fig. 1a), we observed strong ASAP3 expression in many neurons in acute cortical slice preparations (Fig. 1b and d). We recorded spontaneous neuronal activity using a RAMP microscope from neurons with high ASAP3 expression levels by setting single recording voxels on the cell somata at a sampling frequency of 2 kHz (Fig. 1c and d). Large (>10%) and rapid AP-like fluorescence spikes (fAPs) were observed in one neuron (Fig. 1c, pink), while a different nearby neuron was silent over the course of the recording (green), indicating that the optical spikes were not due to noise of the system. The strong sparse expression of ASAP3 and RAMP enabled optical readout of fast neuronal electrical activity in single trials even from a single voxel of neuronal soma. We refer to the combination of ASAP3, IUE and RAMP as AIR.

**Figure 1.**
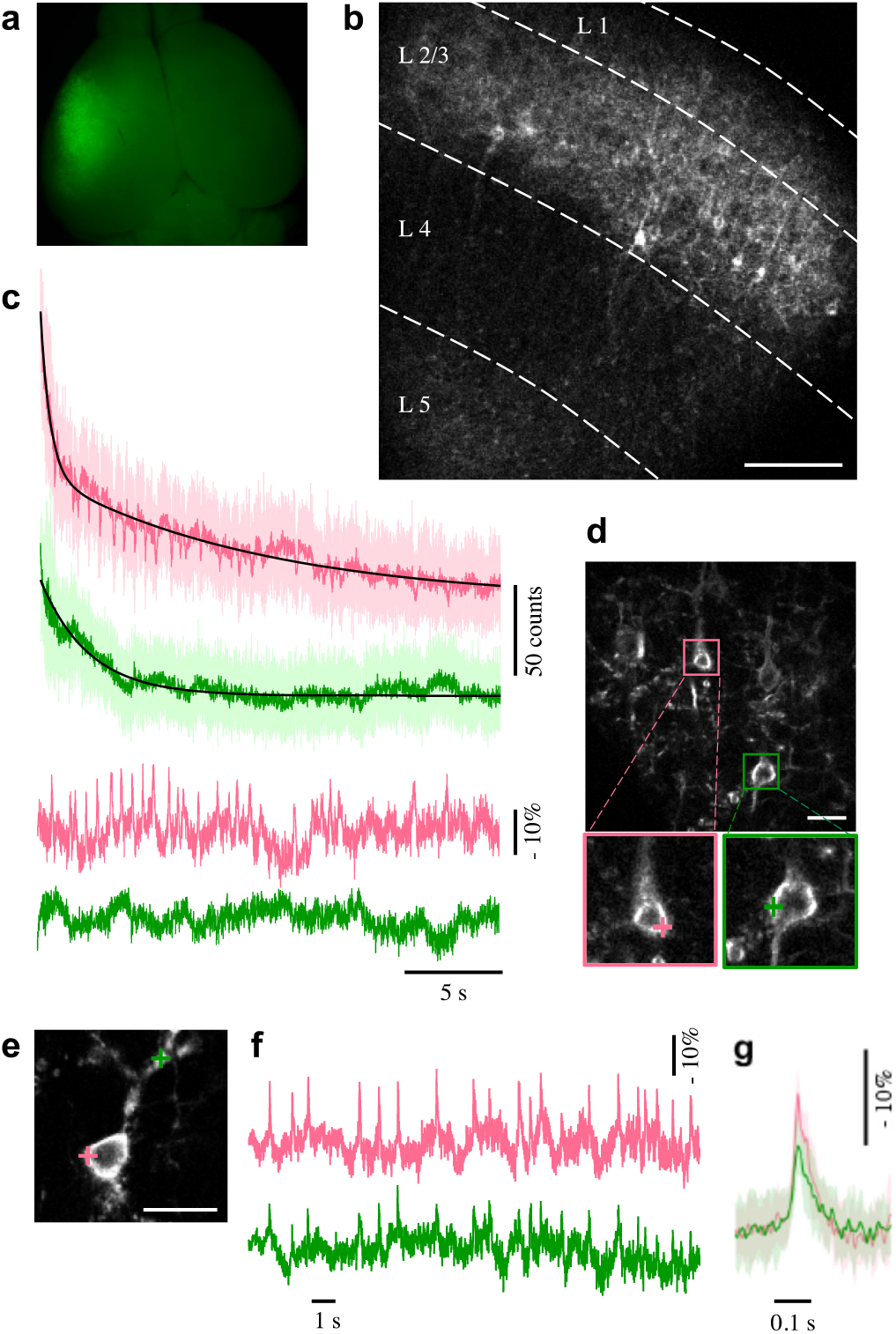
Expression of ASAP3 in murine somatosensory cortex via IUE and membrane potential recording with random-access, single-trial, and single-voxel two-photon voltage recording from multiple sites in acute slice. **(a)** Mouse brain at post-natal day (P) 8 after electroporation of pCAG-ASAP3b at embryonic day (E) 15.5. **(b)** Representative two-photon single-plane image of ASAP3-expressing neurons in an acute cerebral cortical slice imaged by RAMP with low magnification. Several neurons are seen in L2/3. **(c)** Representative single-trial single-voxel voltage recording of spontaneous activity recorded from the two colored neurons simultaneously **in d.** Top, raw (lighter colors) and filtered (50 Hz Butterworth low-pass filter; darker colors) optical traces. Bottom, the traces were corrected for the photobleaching and normalized to the baseline drift (black traces in top). (**d)** Top, a two-photon image of ASAP3-expressing neurons with high magnification. Bottom, magnified views of the boxed regions in top. Colored crosses indicate selected voxels on the plasma membrane for voltage imaging in c. **(e, t)** Representative simultaneous recording of spontaneous somatic (pink) and dendritic (green) activities in a neuron. **(g)** Averaged spike traces taken in the two locations in e. Traces were aligned to onset of somatic activity. The shaded regions denote SD. Scale bar in **b:** 100 µm; in **c** and **e:** 20 µm.

We next evaluated AIR’s ability to detect spontaneous dendritic activity in acute slices. Spontaneous voltage signals were simultaneously recorded from soma and a dendrite of a single cell at a sampling frequency of 10 kHz (Fig. 1e and f). fAPs were observed in the dendrite with perfect synchronization to the somatic fAPs (JBSI = 1.0, Z = 5.39) with a 0.4-ms delay at the peak and reduced peak amplitudes (14.0 ± 2.5% at soma and 9.2 ± 2.9% at dendrite 33 µm from the soma; mean± standard deviation (SD), n = 20 spikes; Fig. 1g). This delay indicates that APs were first generated at or close to the soma and then back propagated to the dendrite. The reduction in the amplitude of fAPs can be attributed to the expected difference in amplitude of APs at soma and in dendrites ^32^. This result indicates that AIR can report fast subcellular voltage changes of dendritic activity as well as in the soma in single trials to study intracellular membrane potential propagation.

We evaluated AIR’s ability to monitor spontaneous activity from multiple neurons simultaneously in acute slice. It requires sufficient SNR for single-trial, single-voxel recording at high sampling rate. In example presented above, optical spikes were easily identified in the fluorescence intensity time course trace recorded at 10 kHz from 2 voxels, where dwell time for each voxel was less than 40 µs. Dwell time is important for good SNR. Thus, to record more neurons simultaneously with sufficient SNR, we reduced the sampling rate to 2 kHz to keep dwell time above tens of microseconds. At 2 kHz, spontaneous optical spikes could be clearly detected from multiple neurons without decrease in SNR. For example, in Fig. 2, the 3 neurons showed synchronized bursts and unsynchronized regular fAPs (Fig. 2b-c), with SNR > 5. In addition, plateau of subthreshold depolarizations (Figs. 2c1, 3a and Supp. Fig. 1) and following hyperpolarizations were observed (Fig. 2f). When the number of recorded neurons was further increased to 6, dwell time was about 72 µs and the SNR was still adequate (> 5) for detecting fAPs and subthreshold membrane potential changes (Supp. Fig. 1). Each fAP within high frequency bursts was clearly resolved, with the shortest interspike interval of 17 ms (Supp. Fig. 2). Taken together, these results indicate that AIR reliably reports fast spiking events and subthreshold events in soma and dendrites from multiple cells simultaneously in brain slices.

**Figure 2.**
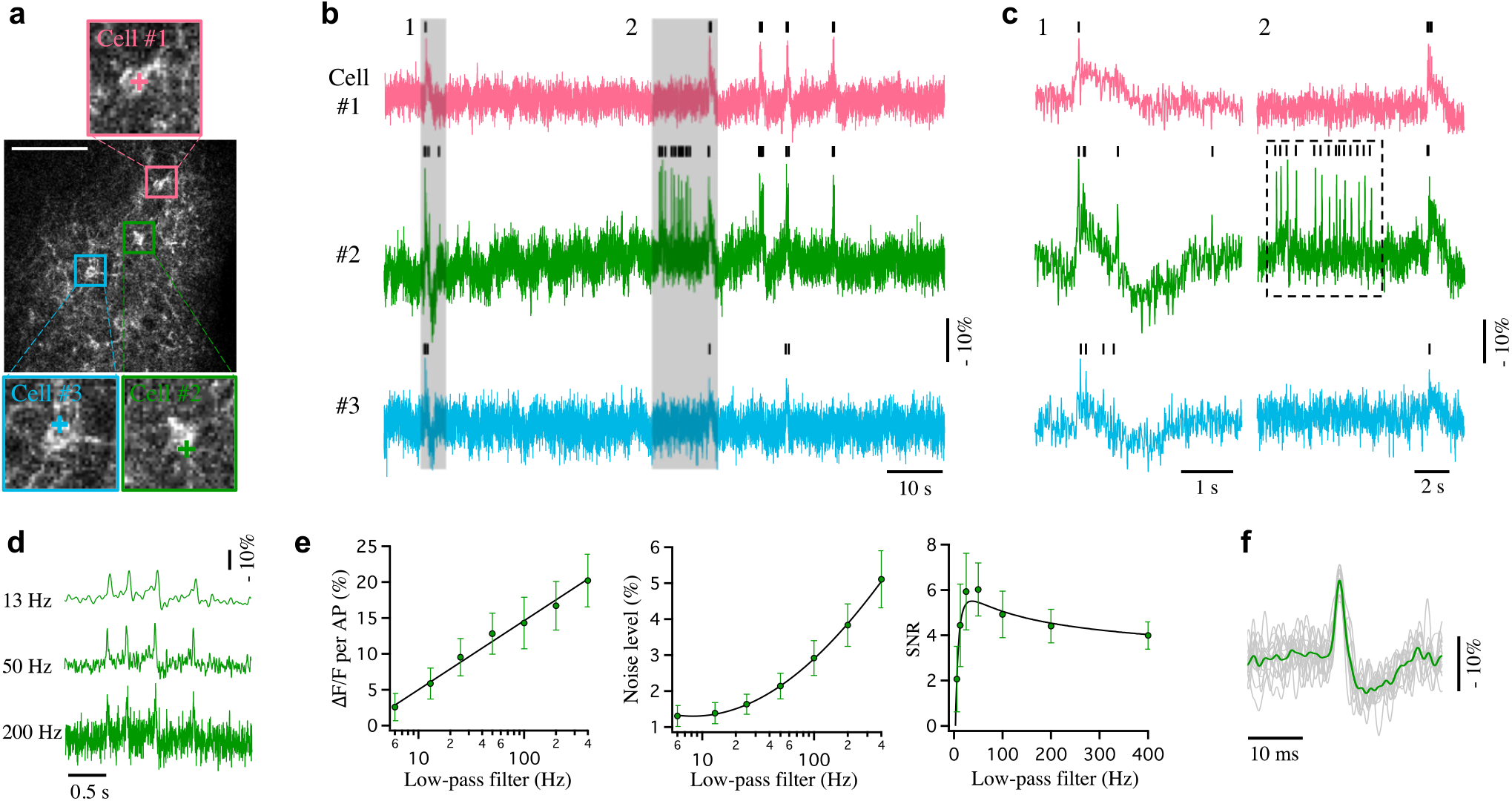
Random-access, single-trial, and single-voxel two-photon voltage recording of multiple neurons and effect of low-pass filter on SNR in regular firing. **(a)** Representative ASAP3-expressing slices imaged by RAMP microscopy in a slice (P9). Regions in colored boxes are magnified to show recorded cells. Colored crosses indicate recording points. Scale bar: 100 µm. **(b)** Spontaneous activities in the color-matched neurons in **a**, recorded at 2 kHz. **(c)** Shaded areas in **b** are expanded, showing clearly distinguishable spikes and subthreshold depolarizations: bursts of fAPs with subthreshold depolarizations and hyperpolarization (**cl)** and regular firing (**c2**, hatched box). Small vertical bars on the top of traces indicate spike events detected. (**d-t)** The spikes in a regular firing in the hatched box in c2 were further analyzed: the raw optical trace was filtered with Butterworth low-pass filters cutting off at 6 - 400 Hz. Three representative traces of 4 spikes are shown in (**d).** The signal level was linearly correlated with the logarithm of filter frequency (*log*_10_(v); **e**, left), and the noise level was correlated with *log*_10_(v) by a power law (n = 14 spikes; e, middle). SNR calculated by dividing the signal value by the noise value had a peak at 37 Hz cut-off (**e**, right). The symbols and error bars indicate mean and SD respectively. The 14 fAPs in the hatched box in **c2** were overlapped **(f**, grey traces), showing their uniform kinetics with overshoot and afterhyperpolarization. Green plot indicates average of the fAPs. The optical traces in **b, c** and **f** were filtered with a 50 Hz Butterworth low-pass filter.

To faithfully detect neuronal activities with AIR high SNR is crucial, therefore we examined low-pass filter parameters as an effective way to reduce noise. Fluorescence trace containing 14 regular fAPs in Fig. 2c2 was filtered with a Butterworth low-pass filter cutting off at 6 −400 Hz (Fig. 2d). The signal level (−ΔF /F) was linearly fitted with the common logarithm of filter frequency (−ΔF/F = *B* + *A* × *log*_10_ (*ν*), A and B are constants, v corresponds to frequency; Fig. 3b, left), and the noise level was well fitted with *log*_10_ (*ν*) by a power law (N/F = *B* + *A* × *log*_10_ (*ν*)^*pow*^; Fig. 3b, middle). The determined power of 0.502 ≈ 0.5 is well in agreement with a prediction.^8^ The relationship between SNR and the filter cut-off frequency (Fig. 2b, right) showed that optimal filtering should occur at 37 Hz to gain the largest SNR for this particular optical recording condition. We evaluated the optimal filter cut-off frequencies in fluorescence traces showing variety of firing patterns, e.g. regular (hatched box in Fig. 2c2 and Fig. 2e) and burst (Fig. 3a and b) firing patterns, obtaining 50.8 ± 13.1 Hz (n = 6 neurons, firing both regular APs and bursts) as the mean optimal filter frequency. Thus, we used a Butterworth low-pass filter cut-off at 50 Hz in this study.

**Figure 3.**
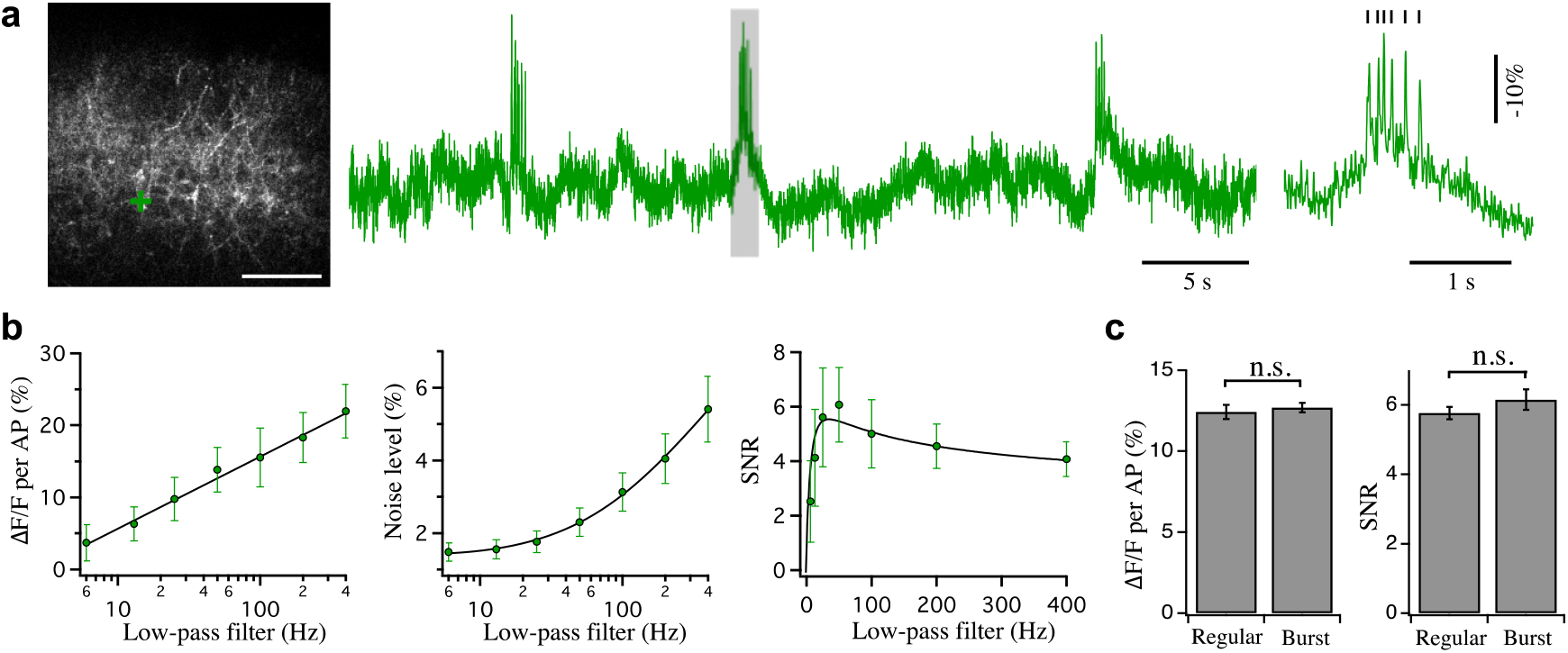
Effect of low-pass filter on SNR in burst firing. **(a)** A representative burst firing in L2/3 neurons in a slice (P9). Left, a bursting neuron (green cross). Scale bar: 100 µm. Middle, a burst firing trace recorded from the marked neuron. Right, expanded view of the shaded area. Fast and narrow spikes are clearly separated. **(b)** A burst spike train was analyzed in the same way as was done in Fig. 2e from the 21 fAPs in the bursts shown in the middle panel of a. The fitting curve (black) in the right graph has maximum SNR at 41 Hz. **(c)** Signal size of fAPs (left) and SNR (right) in regular firing APs (n = 291 spikes in 6 cells) and APs in bursts (n = 214 spikes in 6 cells). Error bars indicate SD.

After filtering, the mean peak amplitude of fAPs (−ΔF /F) was 12.6 ± 0.9% and the SNR was 6.0 ± 0.6% (n = 12 neurons) in single trial recordings from single voxels. We observed no significant differences in peak amplitude and SNR between neuron groups that showed regular APs or AP bursts (Fig. 3c, n = 6 neurons in each group). The filtering resulted in a millisecond-order spike-timing accuracy. Difference in spike timing in traces recorded from two different locations of the same cell body was 1.6 [interquartile ranges (IQR), 1, 2.8] ms, lower than calcium recording with RAMP in barrel cortex slices^33^. Lowering filter cut-off frequency below 50 Hz impacted temporal resolution due to over-filtering. Higher cut-off frequencies caused insufficient filtering with larger noise, resulting in larger temporal errors in peak detection (Fig. S3).

Specific cortical layers can be selected for transfection by simply adjusting the timing ofIUE. Specifically, performing IUE at E13.5 leads to ASAP3 expression in L5 neurons, while IUE done at E14.5 enables voltage recording in L4 and L2/3 neurons. Fig. 4 shows examples of voltage recording in L5 neurons, while Fig. S4 shows voltage recording from L4 and L2/3 neurons. Sequential IUE has also been shown to be effective to express genes in multiple cortical layers ^34^ Thus, AIR permits recording intralaminar and interlaminar neuronal activities.

**Figure 4.**
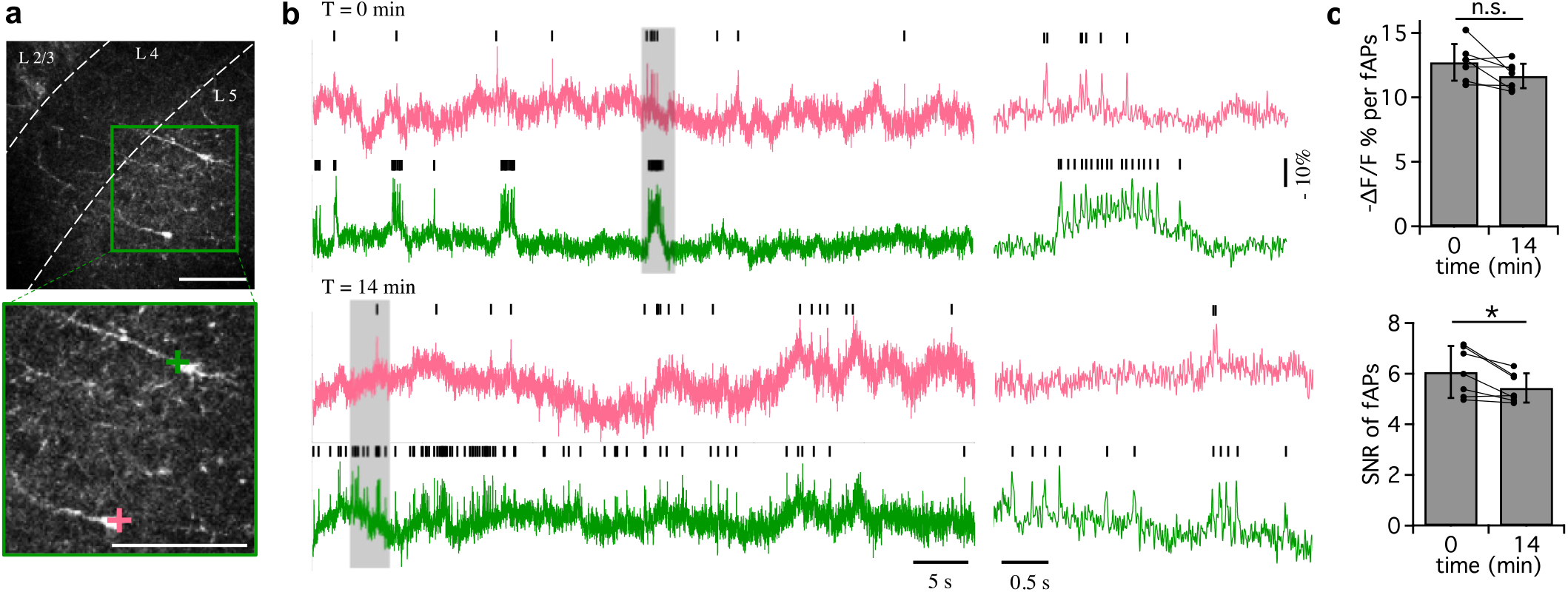
AIR permits long-term recording. **(a)** ASAP3 expressed in L5 neurons in a cortical slice (P8) with IUE at E13.5. The area in the green box in the top image is enlarged in the bottom image. Crosses indicate recording points. Scale bar: 100 µm. **(b)** Optical traces of AIR imaging show the first (top) and last minute (bottom) of a total 14 min recording which was performed by moving recording voxel on the same cell body every 2 min to avoid photobleaching. Shaded areas are expanded in right. Optical spikes are clearly distinguished throughout the recording (small vertical bars). **(c)** Pair-matched comparison of peak amplitude (top) and SNR (bottom) between the optical traces of the first and last minutes. Mean ± SD together with values from each cell are shown.

Next we explored whether AIR could report voltage changes over long time courses. Because SNR is related to the square root of emission photon flux, SNR tended to decrease during long recordings due to photobleaching. A previous ASAP3 study that adopted ultrafast local volume (ULOVE) two-photon excitation reported a 22% photobleaching during the first 3 min of recording ^7^. In this study we observed ~40% decrease in fluorescence during the first 3 min due to the small recording volume. Because photobleaching was limited to the vicinity of the recording spot, the recording voxel was moved to other locations of the same cell body every 2 min to maintain adequate SNR in long duration recordings. This suspended optical recordings for several seconds while the laser spot was moved to a new location. Fig. 4 shows an example of a long-duration recording: membrane potential changes of two neurons were tracked for 14 min in total, and fAPs were easily identified throughout the recording (Fig. 4b). During the recording, the peak amplitude of fAPs tended to reduce by 7.8 ± 3.5% (Fig. 4c; p > 0.05, paired t-test, n = 7 neurons), while SNR reduced by 9.5 ± 2.7% (p < 0.05, paired t-test), but both were still sufficient for reliable peak detection (SNR_14min_ = 5.4 ± 0.6). In one case we were able to monitor a neuron with AIR for 27 min with SNR above 5.

Taking the advantage of the ability to perform long-duration multi-cell recordings, we monitored neuronal activities in circuits with AIR to investigate correlation between neuron pairs in P16-17 somatosensory cortex by a synchrony analysis, in which we adopted a jitter-based method to detect significantly correlated neurons and to quantify the correlation^35^. In an example (Fig. 5), neurons 1, 2 and 3 were correlated with each other, while neurons 4 and 5 fired a lot of spikes but had no significant correlation with other neurons. Out of 106 pairs of L2/3 neurons from 13 mice that had more than 6 fAPs, 46 pairs were judged to be statistically correlated (p < 0.05), with considerable variability in correlation strength (Fig. 6a). Within the significantly correlated neuron pairs, neuron pairs with shorter horizontal distance tended to be more highly correlated (Fig. 6b), which is consistent with previous studies using electrophysiology^29^, photo-stimulation^30^, or calcium imaging ^31^. Neuron pairs with shorter horizontal distance had shorter delay of correlation than pairs with longer distance (Fig. 6c).

**Figure 5.**
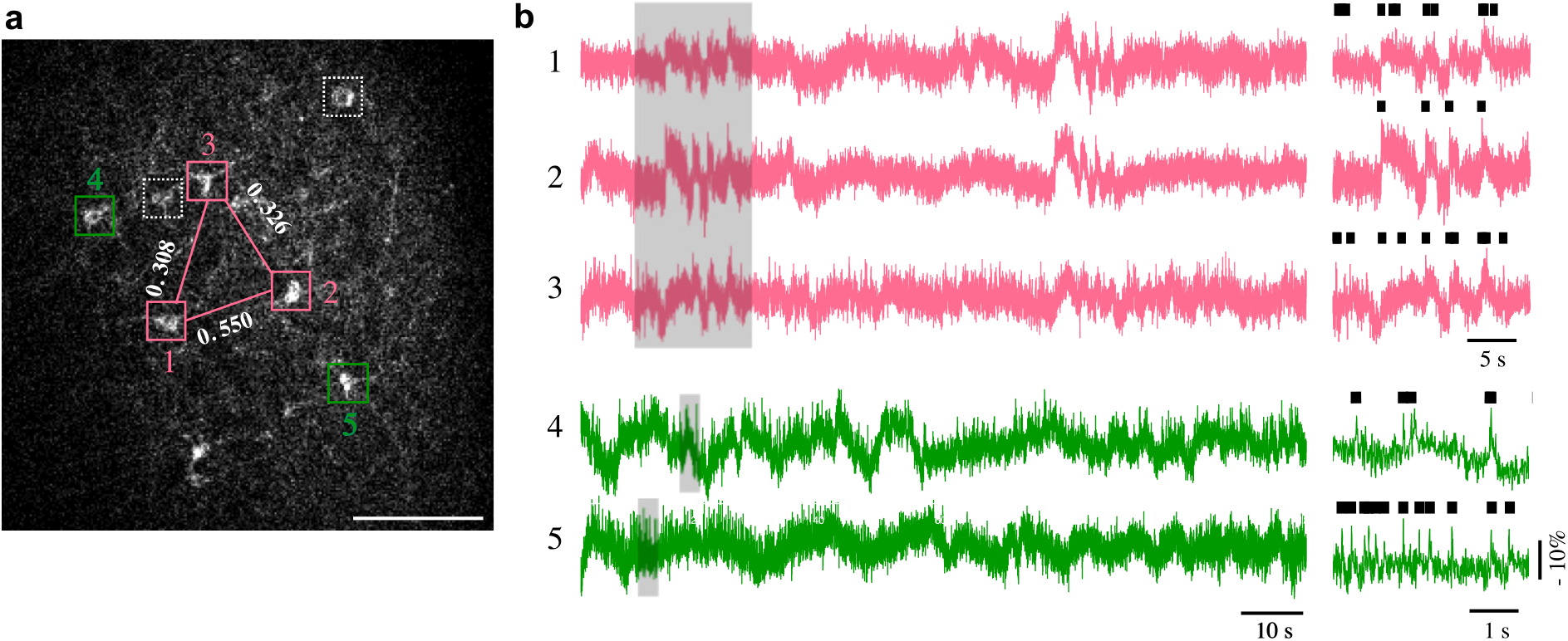
Pairwise correlations of neuron pairs by firing timing **(a)** ASAP3-expressing in L2/3 neurons in a slice (P13). Five neurons showed more than 6 fAPs (colored boxes), and other 2 (white dashed boxes) were silent. Pairwise correlations between each neuron pair was calculated. The 3 neurons in pink boxes were significantly correlated. Strength of pairwise correlation is indicated by white figures. The 2 neurons in green boxes were firing but not significantly correlated with neurons in pink boxes or with each other. Scale bar: 100 µm. **(b)** Spontaneous activities in the neurons displayed in **a.** The 3 pink traces exhibited several synchronized fAPs as well as unsynchronized fAPs, while the 2 green traces only had unsynchronized fAPs. Right, shaded areas are expanded, showing synchronized (pink) and unsynchronized (green) neurons. Detected fAPs are marked by vertical bars.

**Figure 6.**
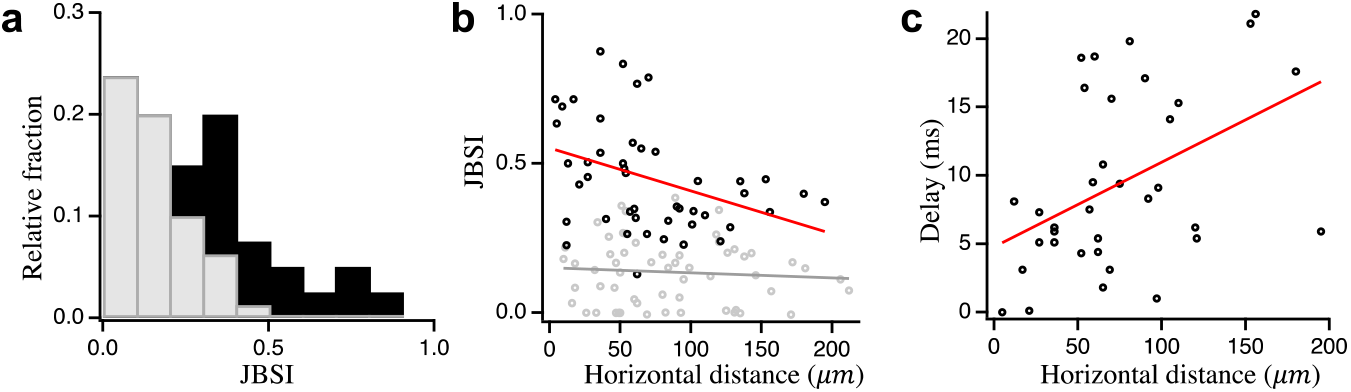
Pairwise correlations of firing neurons **(a)** A histogram showing the fractional occurrence of synchrony index (jitter-based synchrony index, JBSI) of 106 neuron pairs from all mice (P7 - P17). Black bars indicate synchronized pairs (Z score > 1.96), and gray bars indicate not synchronous ones. **(b)** JBSI was negatively correlated with horizontal distance in significantly correlated neuron pairs (black dots) but not in uncorrelated pairs (grey dots; Pearson’s correlation coefficient= −0.37, p = 0.01; Spearman’s rank correlation: r = −0.39, p = 0.007, and Pearson’s correlation coefficient= −0.08, p = 0.55; Spearman’s rank correlation: r = −0.05, p = 0.68, respectively). Red and grey lines indicate linear regression of the synchronous and asynchronous pairs, respectively. **(c)** A positive correlation between the delay of correlation and horizontal distance in significantly correlated neuron pairs (35 neuron pairs from 9 mice; Pearson’s correlation coefficient= 0.44, p = 0.007; Spearman’s rank correlation: r = 0.44, P = 0.009).

In somatosensory cortex, neurons in L2/3 have functional intralaminar horizontal connections and are also connected with L4 neurons^28^. However, the postnatal developmental time course of these neuronal connections has not yet been fully understood at cellular resolution. To investigate the development of neuron correlation in L2/3 of somatosensory cortex, we recorded spontaneous activities of multiple neurons simultaneously in acute slices from P7 to P17 mice, and analyzed activity correlations in all neuron pairs: data was taken from all active neurons found in fields of view irrespective of how they correlated with each other (Fig. 7). In slices from the youngest stage (P7-9), L2/3 neurons were rarely correlated (Fig. 7a, left; median JBSI = 0.08 [0, 0.21], n = 27 neuron pairs from 4 animals). The correlation strength of neuron pairs increased at the P10-12 state (0.21 [0.15, 0.32], n = 29 from 3 animals) and further at the P13-15 stage (0.34 [0.26, 0.50], n = 19 from 3 animals). And, in P13 and older slices, the correlation strength remained high. Distribution of the horizontal distance of all imaged neuron pairs did not differ much along the developmental stages (Fig. 7a, right). Therefore, the age-dependent increase in the correlation strength does not reflect a bias in distances between imaged neurons. These results thus suggest that somatosensory cortex L2/3 neurons increase in horizontal connection strength during the second postnatal week. This is consistent with previous electrophysiological studies^36, 37^, with the AIR approach providing higher throughput with better spatial resolution.

**Figure 7.**
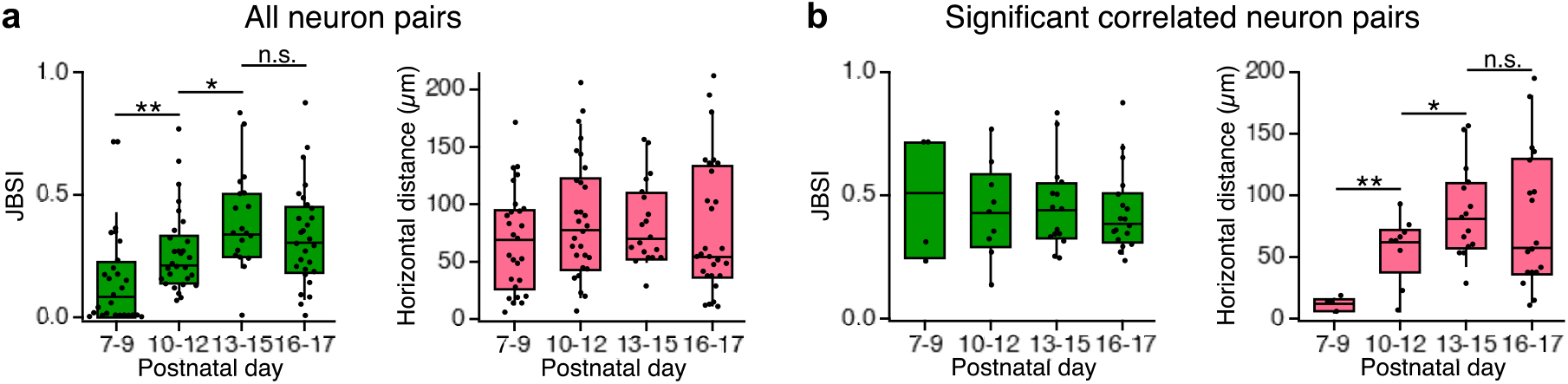
Developmental changes in neuronal correlation in layer 2/3. **(a)** Strength of synchrony (left) and horizontal distance (right) at different postnatal stages in all neuron pairs (n = 27, 29, 19 and 28 pairs for stages P7-9, P10-12, P13-15 and P16-17, respectively). Left, synchrony index in JBSI increased along the developmental stages. Right, there was no obvious difference in horizontal distance of neuron pairs among different stages (p > 0.1). **(b)** Synchrony strength (left) and horizontal distance (right) of significantly correlated pairs at different postnatal stages (n = 4, 9, 15 and 18, respectively). Left, synchrony index was similar among the stages. Right, the horizontal distance of neuron pairs at P7-9 was much shorter than other stages, and that at P10-12 was also shorter than P13-15.

We next asked whether the timing of intralaminar connectivity can be inferred from the time course of synaptic correlation. When neuron pairs with significant correlation were extracted, the correlation strength was within a similar range and did not show a developmental change (Fig. 7b, left). However, the horizontal distance of the correlated pairs was very short at P7-9 (Fig. 7b, right; 12 [10, 13] µm, n = 4), indicating that the L2/3 neurons are synaptically correlated only with adjacent neurons at this stage. In the later stages (P10-12 and P13-15), synaptically correlated L2/3 neurons were located further apart. Thus, the developmental increase in correlation strength when all neuron pairs were tested irrespective of correlation (Fig. 7a, left) is attributed to increased correlation in distant neurons in late stages rather than increased correlation strength in each correlated neuron pair. This result is consistent with previous electrophysiology studies^36^, but with better spatial resolution.

A previous study using the chemical calcium indicator Oregon Green BAPTA (0GB) found that network activity in L2/3 and L4 of somatosensory cortex was more synchronized in postnatal week 1 than in week 2^27^. This seemingly different finding, however, may be due to differences in experimental systems. The previous study recorded neuronal activity *in vivo* with an intact thalamocortical (TC) circuit, which is crucial for synchronized activity in neonatal mice^38^. We desired to address only the question of horizontal connectivity within cortex, and thus used coronal cortical slices in which cortex and subcortical structures are disconnected^39^. Second, our ability to detect and precisely time individual spikes with ASAP3 allows for correlated sparse activity to be detected more efficiently than with 0GB. Specifically, the earlier study reported that calcium imaging accurately detected bursts but missed 60% of single spikes^27^, which is consistent with another study finding 0GB failing to detect at least 75% of active cells in somatosensory cortex under two-photon imaging^40^.

To determine if our ASAP3 traces were consistent with the OGB-based findings, we emulated calcium activity using our recorded spike data by removing different fractions of single spikes and performing normalized correlation coefficient (NCC) analysis as performed in the previous work^27^. When 100% or 60% of single APs were removed, pairwise correlations of neuron pairs were higher in early stages than late stages (Fig. 8a, left 2 panels), opposite to our result using jitter-based synchrony analysis of all spikes (Figs. 7a, left and 8b, left) and matching the findings of Golshani et al. However, when 30% or none of the single APs were removed, correlations of neuron pairs were not apparently different among the stages (Fig. 8a, right 2 panels). Thus, the fidelity of single AP detection impacts the results of pairwise correlation analysis. Our analysis also revealed that a developmental increase in neuron connectivity is only apparent in single APs, because it was diminished by removal of all single APs in jitter-based analysis (Fig. 8b, middle) while it was preserved by removal of all SPs in bursts (Fig. 8b, right). As single spikes became dominant in the late developmental stages than early stages (Fig. 8c), failure of 0GB to detect single APs would result in underestimation of the strength of pairwise connection in late stages. A similar effect may explain the weak relationship between horizontal distance and OGB-based correlation at P13 and later^27^, compared to our detection of distance-dependent correlation at these stages (Fig. 6b). Furthermore, cross-correlation analysis is sensitive to the difference in firing rates between neuron pairs and also to the change in the firing rate^35, 41^. In strongly bursting neuron pairs, the normalized correlation coefficient method tends to overestimate synchrony strength. As neurons in early developmental stage tend to fire in bursts (Fig. 8c), the overestimation of NCC would be more overt in early developmental stage than older stages. Our ASAP3 recording resolved not only APs in bursts but also single APs with high fidelity, and the jitter-based synchrony analysis is not affected by the firing rate difference between neurons and alterations in firing rate. Thus, the ability to detect single APs and obtain their precise timings provides additional sensitivity for functional connectivity analysis.

**Figure 8.**
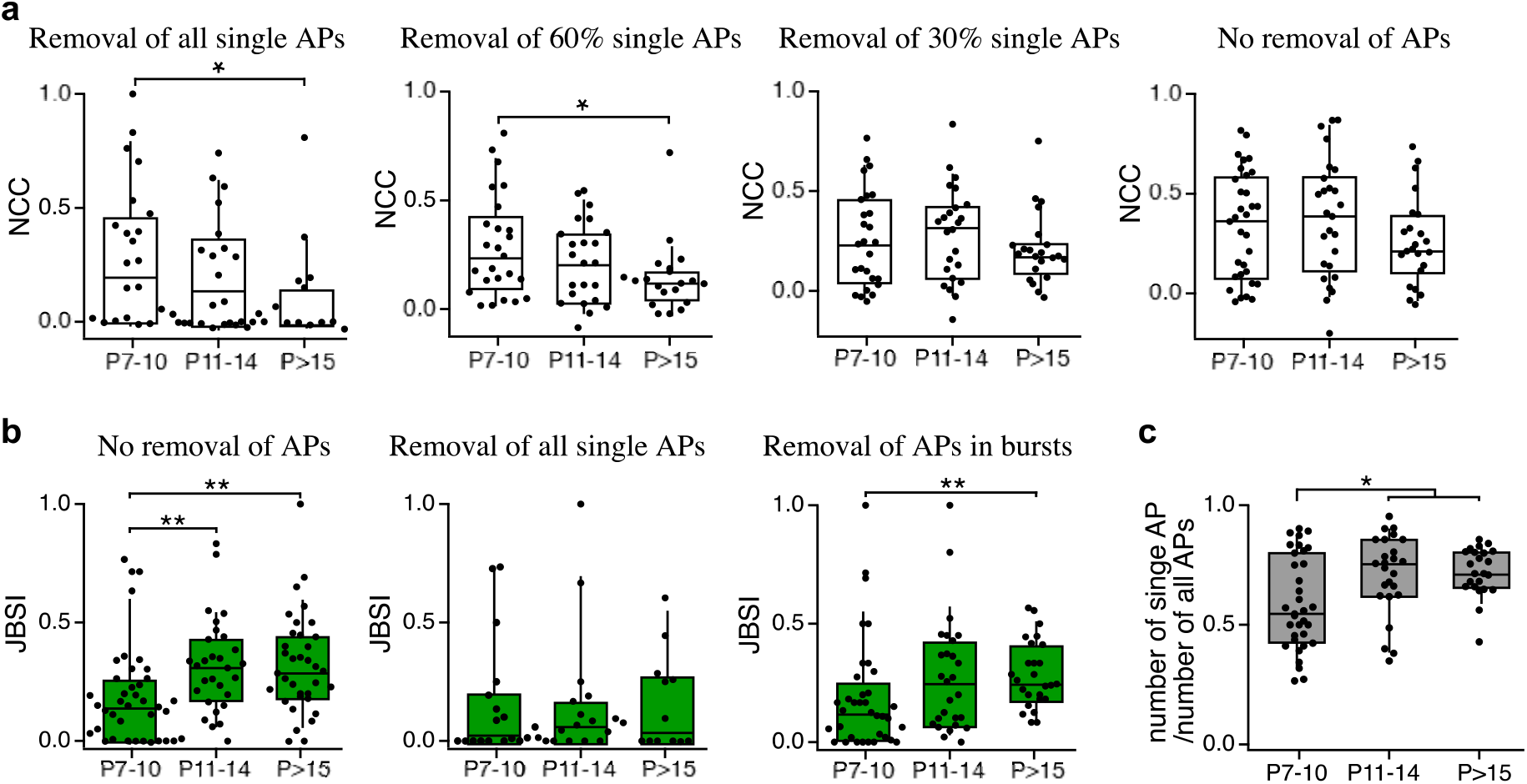
Emulation of neuron connection analysis with calcium imaging by thinning out single spikes. **(a)** Normalized correlation coefficient (NCC) of spike events of neuron pairs were calculated from the same data set used in Fig. 7a after thinning out all, 60%, 30%, or 0% of the single spikes. In the data sets in which all and 60% of single spikes were removed, the strength of pairwise neuron connection showed decrease along development. **(b)** To compare with the NCC analysis method, jitter-based analysis was performed on the same data set without removal of single spikes (left), after removal of all single spikes (middle), or removal of all spikes in bursts (right). The analysis in b, left is essentially the same as the left panel of Fig. 7a but with a different age classification. The "no removal" and "removal of spikes in bursts" data sets showed increase in the strength of pairwise connection along development. **(c)** Ratio of number of single APs to that of all APs in different stages. Black dots represent pairs of neurons. *: p < 0.1; **: p < 0.01, Mann-Whitney U-test.

Taken together, our results demonstrate the ability the AIR approach to characterize spontaneous neuronal activity correlations with greater throughput than electrophysiology and greater accuracy than calcium imaging.

## Discussion

Through continuing development of voltage indicators and advances in microscopy methods, fast voltage imaging is increasingly coming to the forefront of neuroscience as a powerful tool complementary to calcium imaging and electrophysiology in dissecting neuronal circuits. Voltage imaging is easier to perform than electrophysiology, and it offers better temporal resolution than calcium imaging. Although widely anticipated, voltage imaging is less widespread than calcium imaging due to intrinsic constraints related to indicator targeting to the membrane and the fast nature of electrical signals. To overcome these limitations, we utilized a recently developed high-performance GEVI, ASAP3, and achieved high indicator expression in the mouse brain via IUE. To record ASAP3 signals, we used RAMP, which enables multiple cell recording with high sampling rates (2-10 kHz) by selectively scanning only locations containing the indicator. The combination of ASAP3, IUE and RAMP enabled simultaneous monitoring of electrical activity from multiple neurons in brain tissue over the course of tens of minutes. This permitted analysis of neuronal connectivity with better throughput than previously possible.

While IUE provides strong expression early in development, useful for this study, expression is reduced during growth of the animal because the transgenes are not integrated into genome. This drawback of IUE can be overcome with transposon expression system, such as Tol2 ^42^ and piggyBac^43^ by inserting the transgenes into the chromosome. PiggyBac transposon with IUE has been shown to be effective in long-term labeling in cerebral cortex^44^. We confirmed that PiggyBac transposon with IUE offers sufficient expressing level of ASAP3 in adult mice (Supp. Fig. 5). Thus, IUE may be a useful method for long-term expression of indicators in the brain, bypassing the need for intracranial viral injections in some cases.

Adding a Kv2.1 proximal retention and clustering segment to the C-terminus of GEVI results in enrichment of the indicator protein at the soma.^7, 9, 12, 44^. This soma-targeting expression greatly reduces mixing signals from different cells, which is more crucial for one-photon voltage imaging but less for two-photon imaging.

Although cross-correlation analysis has been widely used in determining neuron activity synchronicity, many cross-correlation analysis methods that require spike trains following Poisson distribution have serious limitations when used in real spike trains. In this study, we used a Jitter-based method to compute the correlation between neurons with a well-normalized index, JBSI, which is applicable to any spike trains, without assuming specific firing statistics^35^. Using JBSI, which is not sensitive to mean firing rate or changes in local firing rate, we were able to compare strength of correlation among neuron pairs from different experiments. Although we only focused on excitatory correlations such as synaptic connections and common inputs, further analyses on inhibitory correlation and non-spiking activity would provide more information about neuronal circuits.

A negative correlation between horizontal distances of neuron pairs in L2/3 and their correlation strength after postnatal development had been observed in previous studies using electrophysiology^29^, photostimulation^30^, or calcium imaging^31^. Using ASAP3, we observed a similar negative correlation in P16-17 somatosensory cortex (Fig. 6b). We then further investigated how pairwise neuron correlations in L2/3 of somatosensory cortex change during development. Previous studies have reported that P10-14 is critical for vertical connections between L4 to L2/3^37, 45^ and P13-16 is critical for horizontal connections in L2/3^36^ with electrophysiological methods such as field-stimulation and/or field-recording. In this study, we observed an increase in pairwise correlations between the P7-9 and P13-15 periods (Fig. 7a, left). Thus, this result raises additional evidence for a critical period in L2/3 connections.

In summary, we have demonstrated that the combination of ASAP3, IUE and RAMP was able to report voltage transients from multiple neurons in brain tissue at single-trial and at the single-voxel level, with a subcellular spatial resolution and millisecond-scale temporal resolution. In addition, we have shown that the AIR approach enables tracking spontaneous voltage dynamics, not only APs but also subthreshold events, in soma and dendrites simultaneously from multiple neurons. Long-duration recordings from multiple cells revealed spontaneous activity in somatosensory cortical L2/3 neurons and development of horizontal neuronal connection around the end of second postnatal week. We anticipate that the AIR approach will facilitate studies on the structure and function of neuronal circuits.

## Methods

### Animals

Pregnant ICR mice were purchased from Japan SLC, Inc. (Hamamatsu, Japan). The day of vaginal plug observation was designated as E0.5. E13.5 to E15.5 pregnant mice were used for IUE. All animal experiments were performed under the control of the Institutional Animal Care and Use Committee of Waseda University. The experimental protocols for the animal experiments was approved by the Committee on the Ethics of Animal Experiments of Waseda University.

### Plasmid

The pCAG-ASAP3b plasmid^7^ was purified using an endotoxin-free plasmid purification kit (NucleoBond Xtra Maxi EF, Mancherey-nagel, Düren, Germany) according to the manufacturer’s protocol. The purified plasmid was diluted with HEPES-buffered saline (HBS, in mM, 20 HEPES, 115 NaCl, 5.4 KCl, 1 MgCl_2_, 2 CaCl_2_, 10 glucose, pH 7.4) to a final concentration of 4-6 µg/µl, and Fast Green solution (Sigma-Aldrich, Tokyo, Japan) was added at 0.01% to the solution to monitor the injection.

### JUE

IUE was performed at E13.5, E14.5 and E15.5 for ASAP3 expression in cortical layers 5, 4 and 2/3, respectively^46^. Briefly, pregnant mice were deeply anesthetized by an intraperitoneal injection of sodium pentobarbital (Kyoritsu Seiyaku Corp., Tokyo, Japan) at 50 µg per gram of body weight. After the midline incision was made in abdominal skin and wall, one uterine horn was carefully drawn out. Plasmid solution (4.5-5.5 µg/µl, 1-2 µl) was injected into the lateral ventricle of the intrauterine embryos, and then electric pulses (33-35 V, 50 ms, 4 times) were delivered using an electroporator (CUY21, Nepa Gene, Chiba, Japan) with a forceps-type electrode (CUY650P5, Nepa Gene). After that, the uterine horn was returned to the abdominal cavity, and the same procedure was performed on uterine horn. Finally, the abdominal cavity was closed with sutures to allow the embryos to continue normal development. In this study, more than 90% embryos survived after delivery, and the plasmid was successfully transfected into more than 90% brains with strong expression.

### Brain slice

The ASAP3-transfected mice were sacrificed at P7-P21 for slicing. Brains were sectioned coronally at 400 µm thickness in cooled cutting solution containing (in mM): 120 Choline-Cl, 26 NaHCO_3_, 8 MgCl_2_, 3 KCl, 1.25 NaH_2_PO_4_, and 20 glucose with a vibratome-type slicer (LinearSlicer Pro 7, Dosaka EM, Kyoto, Japan), and left to recover in low-K^+^ - type artificial cerebrospinal fluid (ACSF, containing (in mM): NaCl 125, KCl 2.5, NaH_2_PO_4_ 1.25, NaHCO_3_ 26, MgCl_2_ 1, CaCl_2_ 2 and glucose 20, 310-315 mOsm, equilibrated with 95% O_2_/5% CO_2_) at room temperature (RT; 20-22 °C) for at least 1 h before transfer to a recording chamber. The slices were recorded within 5 hours after recovery. During recording, high-K^+^ - type ACSF (difference from the low-K^+^-type ACSF was 122.5 mM NaCl and 5 mM KCl) was used instead, of which K^+^ concentration is also popular in brain slice experiments.

### Fast random-access two-photon voltage imaging in acute brain slices

A custom-built random-access two-photon microscope was used to perform voltage recording as described^17^. In brief, a femtosecond pulsed Ti:Sapphire laser (output = 600-700 mW, Tsunami, Spectra Physics Japan, Tokyo, Japan) was tuned at 900 nm. The laser beam passed through an acousto-optic modulator (AOM; MTS144-B47A15, AA Opto-Electronic, Orsay, Franc) to pre-compensate for spatial distortions introduced by the AODs, and then deflected by two orthogonal AODs (DTSXY-400-850.950, AA Opto-Electronic) to rapidly and randomly scan the field of view. The laser beam was passed through selectable high or low magnification light paths. The laser beam was focused on the sample using a 20x 1.0-N.A. water-immersion objective (XLUMPlan FL, Olympus, Tokyo, Japan) mounted on an upright microscope (BX51WI, Olympus). Emitted photons were collected by photomultiplier tubes (H7422PA-40, Hamamatsu Photonics, Hamamatsu, Japan) in the photon-counting mode after passing through a 500-550nm band-pass filter. The time to move the laser beam between voxels in the field of view was 11.5 µs, regardless of the distance between the voxels. The sampling rate was set to 2-10 kHz. Estimated dimensions of laser spot was 3 µm for the axial axis with radius of 0.35/1.4 µm for high/low magnification ^47^. All devices were controlled by TI Workbench, a custom-made software written by T.I ^48^.

Acute cerebral slices were visualized in a chamber continuously superfused with the high-K^+^ -type ACSF at RT under the two-photon microscope. ASAP3 expressing neurons were identified by two-photon excitation, and voltage recording was performed on bright neurons in the somatosensory cortex. In each neuron, recording points were chosen among bright points on the plasma membrane. Before starting the recording, ACSF was warmed to 33-35°C as ASAP3 has approximately 4-fold faster kinetics at 33-35°C than at RT^7^. For multi-cell recording, voltage recording was performed at 2 kHz with the low magnification light path. When recording spike propagation within neurons, voltage recording was performed at 10 kHz with the high magnification light path to increase spatiotemporal resolution. All devices were controlled and data were acquired with the TI Workbench software running on a Mac computer^48^.

### Data analysis

For each optical trace, high-frequency noise was removed by low-pass Butterworth filter, and periods with shaky baseline were discarded. Then, APs were detected as fluorescence spikes in TI Workbench using a Schmitt trigger method ^17^. The time series of low-pass filtered raw fluorescence intensity, F_raw_(i), had baseline drift (see Fig. 1 c) due to photobleaching, which was corrected by calculating baseline by linear regression with a 100 ms temporal window just before each data point, F_base_(i). Time series of corrected fluorescence intensity change for each point, ΔF(i), was calculated as: ΔF(i) = -[F_raw_(i) - F_base_(i)]. ΔF is positive in depolarization. If ΔF exceeded 4 times of standard deviation (SD) of the entire baseline-corrected data points and kept exceeding SD for more than 2 ms, this data range was determined as a spike event (fAP) with peak amplitude and peak time as the maximum ΔF value within the period and time of the maximum ΔF, respectively. Noise level was defined as SD / F_base_(i), and SNR was calculated as SNR(i) = ΔF(i) / SD.

To determine the spike timing error of the optical recording, fluorescence signals were recorded at two locations of the same soma simultaneously, and the spike timing errors were determined as the peak time difference between the synchronized spikes recorded from the two locations.

Pairwise correlation of neuron pairs was evaluated by a jitter-based synchrony analysis ^35^. In particular, Jitter-based Z-score was calculated from each pair of spike trains derived from multi-cell recordings of spontaneous activity, determining whether the correlation of two spike trains was statistically significant. The Jitter-based Z-score was calculated as:

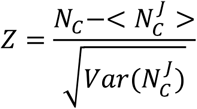

*N*_*C*_ is the number of spike coincidences within a synchrony span, τ_s_. 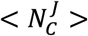 is the expected value of the number of spike coincidences after shifting each spike in the lower firing cell within the jitter span, τ_J_ (see figure 1 of Agmon et al.^35^). And 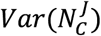 is the variance of the number of spike coincidence after applying the jitter. To quantify correlation strength, jitter-based synchrony index (JBSI) was calculated as:

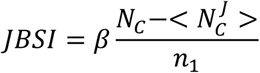

where *n*_1_ is the number of spikes in the lower firing cell of the pair. *β* = 2 when τ_s_ / τ_J_ ≤ 2, and *β* = τ_j_/(τ_j_-τ_s_) when τ_j_ / τ_s_ > 2. We chose τ_j_/τ_s_ to be 2 for simple calculation. Thus, *β* = 2 in our analysis. Neurons showing less than 6 spikes during the entire recording period were discarded. As JBSI is a well normalized synchrony index, which is independent of firing rate, not sensitive to the firing rate difference of neuron pair and not requiring specific firing patterns, it allows valid comparison of correlation strength of neuron pairs obtained from different experiments. JBSI = 1, −1 and 0 indicate perfect positivity synchrony, perfect negative synchrony, and chance-level synchrony, respectively.

Both Z-score and JBSI depend on τ_s_. The slope of JBSI versus τ_s_ curve is gentle on the right of the peak and steep on the left. JBSI has a maximum value when τ_s_ equals to the temporal precision of fAPs, which depends on the lag of spike pairs together with the temporal error in peak detection. Thus, the points at which the JBSI-τ_s_ curve drops off steeply to the left should be used as cutoff τ_s_ for determining spike lag and JBSI calculation. As different neuron pairs may have different spike lags, neuron pairs have their maximum JBSI values (JBSI_max_) with different cutoff τ_s_. JBSI values were calculated while τ_s_ was changed from 1 to 100 ms with a 1 ms step, and the maximum value was determined as JBSI_max_ and corresponding τ_s_ was used as cutoff τ_s_ for each neuron pair. And the τ_s_ value was also used to calculate Z-score. Neuron pairs with Z > 1.96 (corresponding to p < 0.05 for a two-tailed Z-test) were considered to have statistically significant connection. And the value of cutoff τ_s_ was considered to be the delay of the spike synchrony.

To emulate pairwise correlation analysis with Ca imaging, we applied normalized correlation coefficient (NCC) analysis^27, 41^ to the spike time series data set obtained in this study. Before NCC analysis, all or part of single spikes were thinned out from spike time series of each neuron, then binned with 1 s intervals: fAPs with less than 300 ms interval with nearby fAPs were defined as APs in bursts, and other APs were defined as single spikes. NCC of each pair of processed spike time series was calculated using the formula:

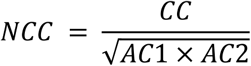

where CC is the cross-correlation coefficient of the spike time series pair (trace 1 and 2), andACl andAC2 are the autocorrelation of time series 1 and 2. When part of single spikes were thinned out, predetermined number of single spikes were randomly chosen and NCC was calculated after removal of the chosen single APs. This random selection and NCC calculation was performed 100 times and average of the NCC values was obtained. Neurons showing less than 6 spikes during the entire recording period were discarded.

Neuron pairs that had more than 6 synchronous spikes were used for analysis of correlation lag. For each pair, correlation lag was determined as the mean of time difference between synchronous spikes.

Data following a normal distribution were represented as means ± SD with bar plots. When normal distribution cannot be assumed, results were represented as median and IQR [25^th^ and 75^th^ percentile] with box plots. Before performing statistical comparisons, the Shapiro-Wilk method was used to test whether the data followed a normal distribution. If so, statistical comparisons between two data sets were performed with two-tailed paired or independent-samples student’s t-test if the two groups had equal variance. When equal variance cannot be assumed, Welch’s t-test was used. If the data do not follow normal distribution, Mann-Whitney U nonparametric test was performed. Statistical correlation between two data sets were calculated with both Pearson’s correlation test and Speannan’s rank correlation test. Statistical tests were performed in Excel (Microsoft) and Igor. JBSI and NCC were computed with custom Python scripts.

For display in the figures, fluorescence intensity traces were filtered and corrected for photobleaching and normalized to baseline using Igor Pro (WaveMetrics, Portland, OR, USA). The procedures were shown in fig. 1c. First, the raw traces (Fig. 1c top, lighter traces) were filtered with 50 Hz Butterworth low-pass filter (Fig. 1c top, darker traces show the filtered data). Then, the filtered traces were fitted with a double or single exponential function (Fig. 1c, top, black traces). Optical traces with initial rapid photobleaching were fitted well with a double exponential function (e.g. the pink trace), and slow photobleaching traces were fitted well with a single exponential function (e.g. the green trace). Finally, the filtered optical traces were corrected for photobleaching and normalized by dividing the traces by the fit function (Fig. 1c, bottom).

## Acknowledgements

This work was supported by Grant-in-Aid for Scientific Research from the Japan Society for the Promotion of Science [JSPS; 23300121 (T.I.)], Private University Research Branding Project (MEXT, Japan, T.I.), and Waseda University Grants for Special Research Projects.

## Author Contributions

All the authors had full access to all the data in this study and take responsibility for the integrity of the data and the accuracy of the data analysis. T.I. and B.L. conceived and designed the research; B.L. preformed the research and analyzed the data; Y.K. contributed unpublished studies; B.L. learned the IUE protocol from S.Y. and K.N. at K.N.’s laboratory in Keio University School of Medicine; M.C. and M.Z.L. created and provided ASAP3; B.L and T.I. drafted the manuscript; M.C. and M.Z.L. provided critical revision of the manuscript; T.I. obtained funding.

## Data availability

The data that support the findings of this study are available from the corresponding author upon reasonable request.

## Code availability

The custom Python scripts are available from the corresponding author upon request. The custom software, TI Workbench, is available on the web site (http://inouelab.biomed.sci.waseda.ac.jp/inouelab-web/tiwb.html).

