## supplemental figures for "Two-photon voltage imaging of spontaneous activity from multiple neurons reveals network activity in brain tissue"

| | |
|---|---|
| Supp. Fig. 1 | Simultaneous voltage measurement in 6 neurons |
| Supp. Fig. 2 | Fast AP trains resolved by AIR |
| Supp. Fig. 3 | Effect of the filter cut-off on timing errors |
| Supp. Fig. 4 | AIR imaging can monitor subthreshold activity in single-trial and single-voxel |
| Supp. Fig. 5 | Imaging voltage with ASAP3 expressed by piggyBac transposon and IUE |

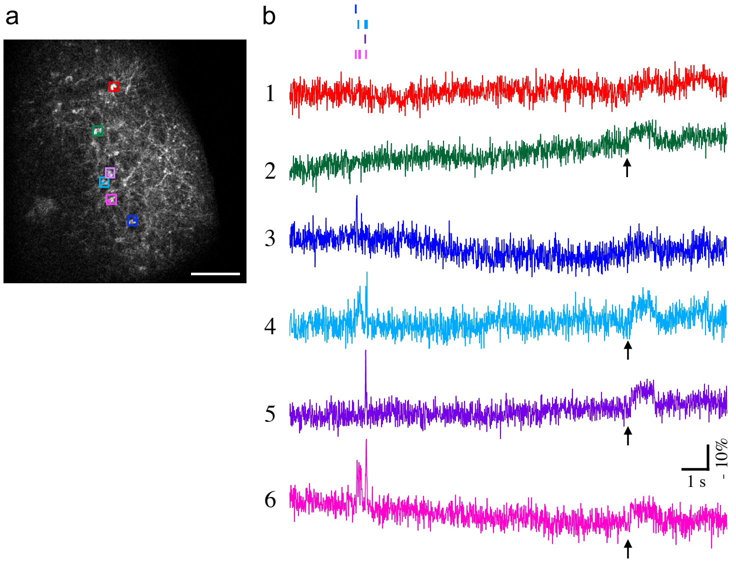


***Supplementary Figure 1****. Simultaneous voltage measurement in 6 neurons*

(**a**) Two-photon image of a cortical slice. Six neurons (colored boxes) that have strong ASAP3 expression were recorded simultaneously. Scale bar: 100 µm. (**b**) Color-matched optical traces recorded from the 6 neurons in **a** at 2 kHz with dwell time 72 µs. Color bars on the top indicate detected spikes in the color matched traces. The 6 traces share synchronized activities in some parts but independent in other parts, indicating that the signal changes are not reflecting noise. In particular, trace 1 is relatively silent; traces 3/4/6 and traces 4/5/6 have synchronized fAPs; and traces 2/4/5/6 show clearly distinguishable plateaus of subthreshold depolarization (arrow).

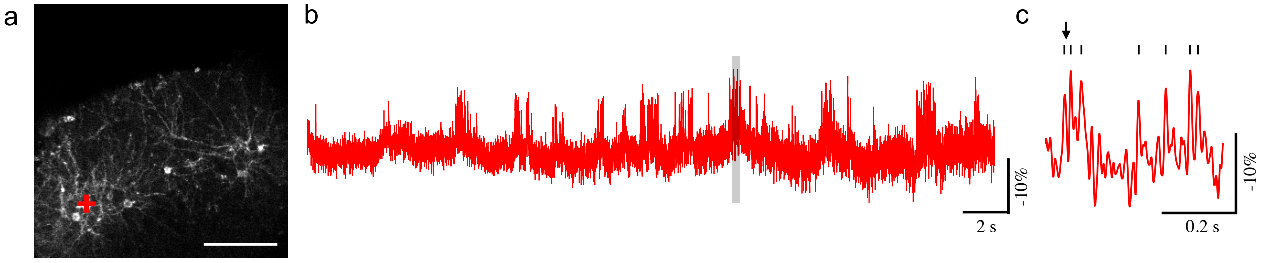


***Supplementary Figure 2***. *Fast AP trains resolved by AIR*

(**a**) An image showing ASAP3 expressing neurons in 2/3 L in somatosensory cortex in brain slice at P16. Scale bar: 100 µm. (**b** and **c**) Fluorescence time course taken from a neuron (red cross in A) at 2 kHz is shown (**b**), and a part of it (shaded area) is expanded (**c**). Spontaneous fAPs were clearly detected (small vertical bars in **c**), and the shortest interval between fAPs was 17 ms (arrow). The optical trace was filtered with a 50 Hz Butterworth low-pass filter.

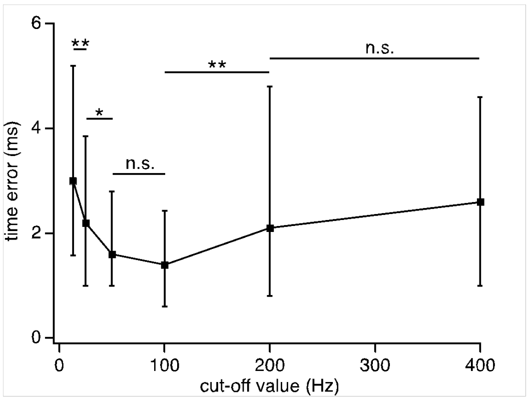


***Supplementary Figure 3****. Effect of the filter cut-off frequency on peak-time detection*

Errors in peak-time detection were determined as the differences in peak time between pairs of optical spikes recorded at two locations of the same soma simultaneously. At filter cut-off frequency of 50 Hz the time error was 1.6 [1, 2.8], which had no significant difference with time error at 100 Hz. While, at lower and higher cut-off frequencies, the time errors were significantly larger than those of 50 and 100 Hz. Square markers and bars represent median and inter-quartile range. *: $p < 0.1$; **: $p < 0.01$, Mann-Whitney U-test corrected with Holm-Bonferroni method for multiple comparisons (n = 4 neurons, containing 152 spikes for each group).

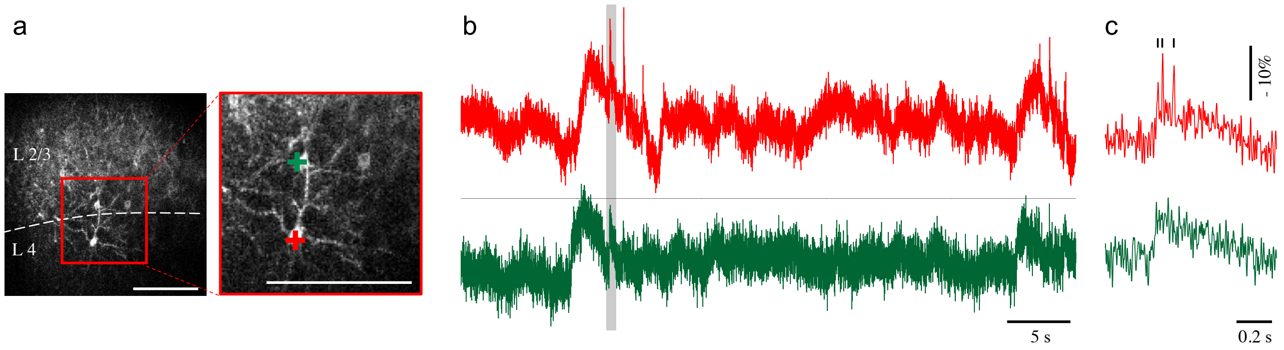


***Supplementary Figure 4***. *AIR imaging can monitor subthreshold activity in single-trial and single-voxel*

(**a**) ASAP3 expressed in L4 and L2/3 neurons with IUE at E14.5. In the bottom image, boxed area in the top image is expanded, showing target pyramidal neurons in L2/3 (green) and L4 (red). Recording points are indicated by crosses. (**b**) AIR recording from the neurons shown in A. The green trace exhibits non-spike events, synchronized with the fAPs in red trace. (**c**) expanded traces of the shaded areas in a. Scale bar: 100 µm.

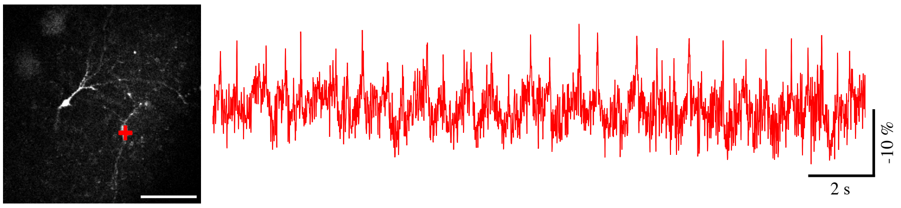


***Supplementary Figure 5****. Imaging voltage with ASAP3 expressed with piggyBac transposon and IUE*

IUE targeting hippocampus was performed at E14.5, and hippocampal slices were prepared at P20. Two pyramidal neurons were identified by the shape and location. Multiple optical spikes were observed in one of them (red cross and trace). Scale bar: 100 µm.
